# Combiroc: when ‘less is more’ in bulk and single cell marker signatures

**DOI:** 10.1101/2022.01.17.476603

**Authors:** I. Ferrari, S. Mazzara, M. Crosti, S. Abrignani, R. Grifantini, M. Bombaci, R.L. Rossi

## Abstract

Here we present the *combiroc* R package, for signatures refinement in high throughput omics. Based on a ROC-driven marker selection, it can be used to find powerful smaller sub-signatures from scRNAseq experiments and to annotate cells using fewer markers. Trained on PBMC dataset, combiroc found NK marker combinations with high cell-discriminating power, in agreement with human protein atlas and that were validated both computationally and experimentally on independent datasets.

## Context and Results

In life science research, smaller number of molecular markers translates into easier annotation and detection, as well as lower cost per testing when applied to diagnostics. Omics methods generate long gene signatures often without sufficient discriminatory power due to carried-over noise. We previously described a R-Shiny application^1,2^ used to select subsets of biomarkers from relatively small signatures^3,4^: this approach was computationally limited and it was mainly applied to diagnostics. Here we expand the method with the new *combiroc* package^5^ that fits more research needs, allowing full customized analysis of larger marker signatures, as they are usually generated by RNA-seq and single cell RNA-seq experiments. This package introduces data importing from established scRNA-seq file formats, automatic detection of signal thresholds and cell annotation features. Using *combiroc* package to well-characterized scRNA-seq datasets, we showed that combinations of up to five individual markers selected from much larger, traditional gene signatures greatly improve the ability to discriminate between different cell type clusters, regardless of differential expression ranking of individual genes in the original signature.

In the context of omics methods, signatures, i.e. lists of markers characteristically expressed in a specific cell cohort, are made of tens, if not hundreds, of features; this entails a huge number of possible combinations, even for a few tens of genes. In the process of identifying a subset of markers from a given signature the most critical step is the choice of a specific signal threshold which is strictly dependent on the nature of the assay. Setting the positivity of detected markers is easier when data is self-produced and the detection system is known; the same procedure can be challenging, or even impossible, when handling others’ or not well documented data, and is not well established for gene expression experiments at the single cell level. To solve this problem, we improved the workflow (**Fig.1a**) introducing a new feature that automatically evaluates the signals’ distributions from both labeled sample classes, and calculates a threshold value to be used in subsequent computations. Density distributions of signals from both classes are plotted and can be visually inspected, along with the suggested signal threshold value (**Fig. 1b**). This method facilitates an important yet arbitrary process guiding the threshold choice in a data-driven way, and it showed to be consistent with previously reported results on AIH datasets^1^. In single-cell RNA-seq, identifying cell population using known markers’ expression is a lengthy manual process. Methods for the automatic or reference-based classification of cells exist^6^, but established marker genes can be individually unreliable for cell classification due to sparsity of scRNA-seq data^7^, and rare cell types can lack unique markers to discriminate them among cell populations. To address this issue we used *combiroc* to determine marker combinations for a specific class in a training dataset and used the linear regression model from such combinations to identify cells of that class in an unlabeled test dataset; training and test datasets must be of the same nature and with the same set of markers (**Fig. 1c**). Whole workflow’s code is detailed in the package main vignette (**Suppl. Material 1**).

**Fig. 1.**
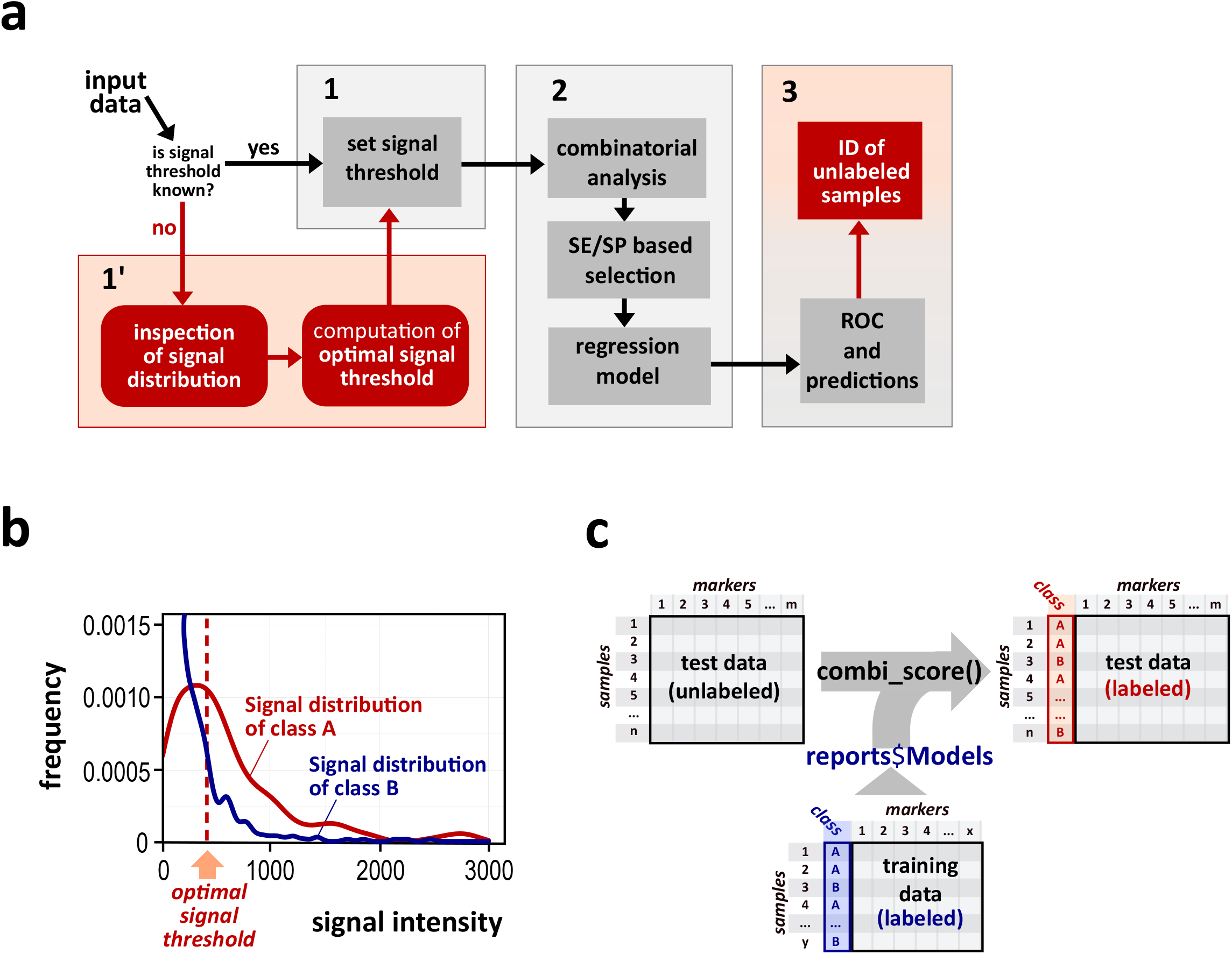
Combiroc automates signal threshold setting and creates models to annotate unlabeled samples. **a**, Breakdown of combiroc’s workflow in phases (1, 1’, 2 and 3). Red boxes highlight new features specifically introduced by the combiroc package: alternative phase 1’ for computation of signal threshold and part of phase 3 with labeling of unknown samples. **b**, Display of the calculated optimal signal threshold (dashed line) on overlapping signal intensity distributions. **c**, Samples of unlabeled data (left) can be associated with a class (right), using regression models generated from labeled training data.

To test if combinations of markers selected by *combiroc* from existing larger signatures could help, we took into account the phenotypic overlap between NK cells and cytotoxic CD8+ T cells^8^ and we assessed the ability of the combinations to discriminate between cell types in the multimodal peripheral blood mononuclear cells (PBMC) CITE-seq atlas^9^, a widely accepted single-cell RNA-seq public reference (**Suppl. Fig. 1a**). Though the NKG7 gene is known as specific to NK cells, in this dataset it is also highly expressed in CD8 T cells (**Suppl. Fig. 1b**); this can lead to an overlapping mislabeling of these two cell types. We determined the NK cells marker genes with the standard Seurat differential expression protocol and took the top 30 genes as the standard NK signature. From this signature we calculated the more than 170-thousand combinations of up to five genes, and we choose to use the best performing ones (ranked by their Youden index) with very high SE and SP values compared to top single markers (**Fig. 2a; Suppl. Fig.2**). To this aim, we tested the models associated with these top combinations on other independent datasets of similar cells: the cord blood mononuclear cells (CBMC) CITE-seq data from the “Seurat multimodal vignette” ^10^; the PBMC-3K scRNA-seq^11^; and the PBMC-Covid19 multiomics datasets^12^ (**Suppl. Fig. 3**). All cell labels were removed from these datasets, only to be used later as ground truth. We assessed the descriptive power of the models in all test datasets calculating the predicted probabilities of being NK cells given a NK cells-specific combination, here defined as “combi-score”. For CBMC dataset the combi-score from comb. #172173 (which has the highest accuracy and it’s made by genes GZMB, IL2RB, KLRF1, SPON2 and TRDC) unambiguously picks the NK cells cluster among the others (**Fig. 2b**). In PBMC-3K dataset we used combination #137550 due to the missing expression of TRDC gene in this dataset: here too, the NK cells were correctly identified (**Fig. 2c**). In the last dataset analyzed (PBMC-Covid19), combiroc correctly identified all types of NK cells (16hi, 56hi, proliferating) that are annotated; moreover, Innate Lymphoid Cells (ILC) were also clearly identified (**Fig. 2d**), which is in agreement with NKs belonging to the ILCs family^13^.

**Fig. 2.**
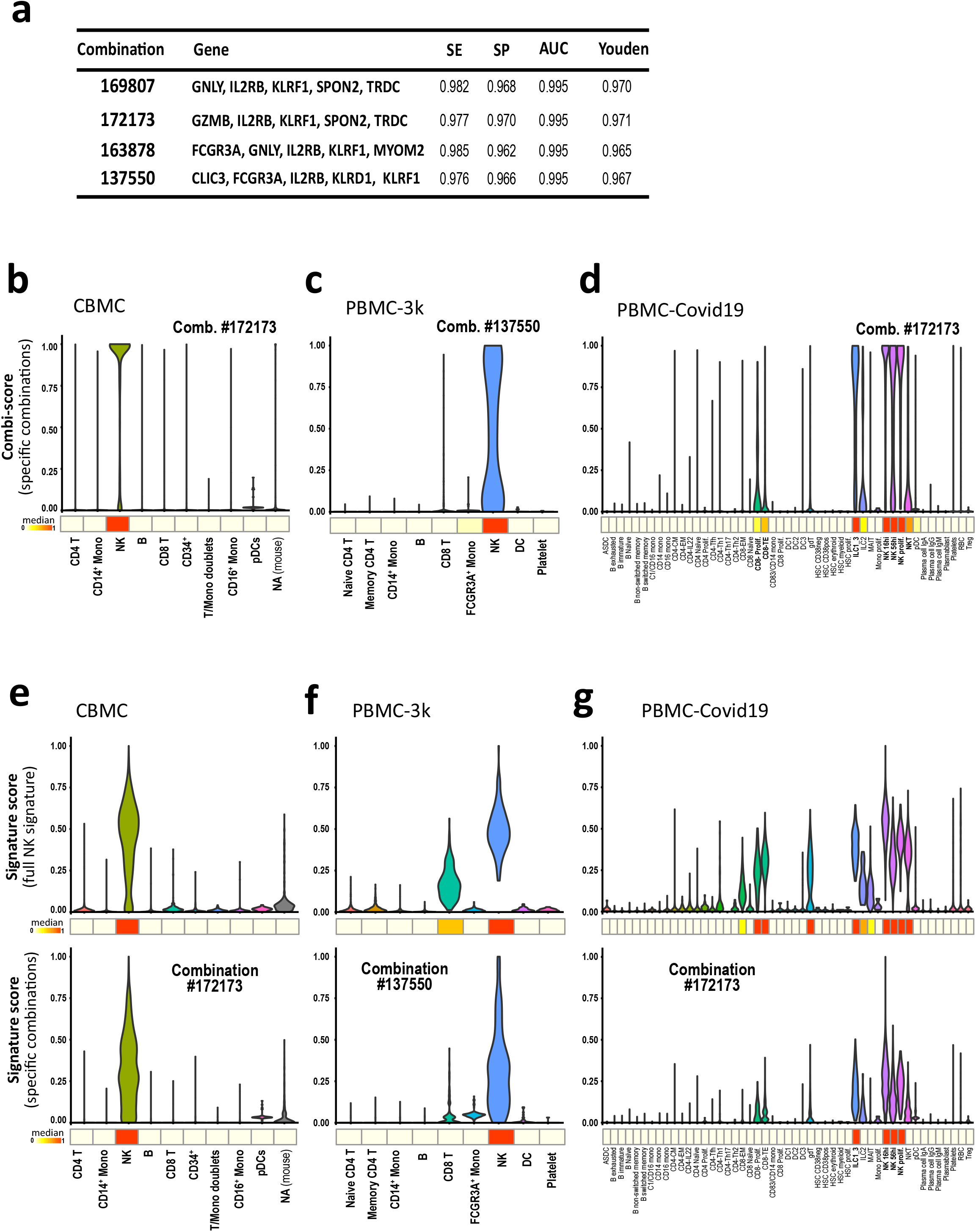
Combiroc combinations correctly distinguish NK cells and describe them more clearly than bigger signatures. **a**, Top four out of 174436 combinations made of 5 markers. Sensitivity (SE), specificity (SP), area under the curve (AUC), and Youden index. **b-d**,Combi-scores (probabilities of being NK cells) computed across cell clusters of test datasets for (**b**) CBMC, (**c**) PBMC-3K and (**d**) PBMC-Covid19 using combination #172172, #137550 and #172173 respectively. **e-g**, Signature-scores from NK cells signature (30 genes, upper panels) compared to signature-scores from combinations (5 genes, lower panels) across the test datasets: CBMC (**e**), PBMC-3K (**f**), PBMC-Covid19 (**g**). The heatmap strips under each panel show the median of the violin plot scores’ distribution

Then we used an independent metric, the “gene signature score” ^14^, to compare the descriptive power of the whole 30-genes NK signature with those of the 5-genes *combiroc* combinations. To do so, we computed the gene-signature scores for the whole 30-genes NK signature in all datasets and compared them to the gene-signature scores for the top *combiroc* 5-genes combinations in all datasets (#172173 in CBMC and PBMC-Covid19, #137550 in PBMC-3K). Remarkably, despite the six-fold reduction (from 30 to 5) in the number of considered features to build these gene scores, the scores derived from combiroc combinations are as accurate in identifying the NK cells as those derived from the whole signature (CBMC) or even better with less noise as in PBMC-3K and PBMC-Covid19: in this last dataset lowering signal from CD8 T cells and gamma-delta T cells, whose median was well below that of NK cells compared to those obtained with the whole signature expression score (**Fig. 2e**). *combiroc*’s marker combinations are thus *de-facto* optimized gene signatures with equal or even higher discriminatory power compared to the larger, parent signature. Code for the single-cell workflow is accessible as a vignette and archive (**Suppl. Material 2-3**).

Finally, we verified that all genes in *combiroc* combinations are coding for proteins that are annotated as enriched in NK cells in the Human Protein Atlas (HPA)^15^ with high specificity scores (**Suppl. Table 1**). NK cells are traditionally sorted using CD56 and CD16 (FCGR3A) levels on CD3 negative cells^16^: since markers coding for membrane proteins can be detected with cytofluorimetric assays, we also evaluated how well the FACS-measured co-expression of a few membrane associated markers from combinations performed on NK characterization compared to standard markers. We found that the combination of cells expressing IL2RB (CD122) and KLRD1 (CD94), in the CD3-negative lymphocyte subpopulation, represents 39% of circulating NK, a fraction similar to that obtained (38.2%) with traditional markers such as CD16+ and CD56dim/bright **(Suppl. Fig. 4a)**. Moreover, CD122 identifies NK cells similarly to conventional methods in terms of cytotoxic activity and cytokine production **(Suppl. Fig. 4b)**. Markers extracted from combinations were thus confirmed with cytometry staining and were expressed on a similar to larger proportion of functionally active NK cells compared to those using conventional markers. In conclusion, we showed that the combi-score can be used to discriminate among cell clusters and that the ability to validate cells’ identity in an unlabelled dataset is very high using a *combiroc*-selected small subset of the original signature. We demonstrated that the few top differential expression genes are not necessarily the most specific ones and that standard DE signatures can be refined with *combiroc*.

## Methods

### Extraction of the NK markers

The 30 genes differential expression signature from NK cells was determined from the Multimodal PBMC dataset from Hao et al. 2021 using the standard Seurat protocol for differential expression with the function FindMarkers(SeuratObject, ident.1 = “NK", ident.2 = NULL), then ordered by Fold Change and the top 30 hits were selected as NK signature.

### Finding the best marker combinations and models

To find the best combinations we used the *combi()* function. This function works on the training dataset by computing the marker combinations and counting their corresponding positive samples for each class (once thresholds are set). A sample, to be considered positive for a given combination, must have a value higher than a given signal threshold (*signalthr*) for at least a given number of markers composing that combination (*combithr*).

As described in the combiroc’s vignette for the standard workflow (**Suppl. Material 1** and GitHub at https://ingmbioinfo.github.io/combiroc/articles/combiroc_vignette_1.html), the argument *signalthr* of the *combi()* function should be set according to the guidelines and characteristics of the methodology used for the analysis or by an accurate inspection of signal intensity distribution. If specific guidelines or knowledge are missing, one should set the value *signalthr* as suggested by the *distr$Density_plot* feature.

The screening of combinations was limited to those composed by no more than five markers (setting the *max_length = 5* in the *combi()* function). While this choice is arbitrary, the reason for this was double: first, it allows a decrease of the number of combinations to compute, making the analysis more manageable; second and more important, it fulfills the original aim of the package which is to trigger easier research and clinical applications, looking for combinations significantly *shorter* than the original gene expression signature. We chose not to set a default for this number since the optimal maximum number of combinations-composing genes can vary depending on the field of application and the experimental and/or clinical context.

### Optimal signal threshold prediction

To predict the optimal signal threshold we used the *markers_distributions()* function, setting the argument *signalthr_prediction = TRUE*. In this way *distr$Density_plot* (see combiroc’s vignette for the standard workflow, **Suppl. Material 1**) will compute the threshold and show it besides the distribution of the signal intensity values for both classes; the threshold is computed as the median of the signal threshold values in *distr$Coord* at which SE and SP are greater or equal to their set minimal values (*min_SE* and *min_SP*). The optimal threshold is added to the “Density_plot” object as a dashed black line and a number, which is being used as *signalthr* value for *combi()* function.

### Training models on selected combination

Regression models on the selected combinations were trained using the function *roc_reports()*, which applies the Generalised Linear Model (*stats::glm()* with argument *family= binomial*) on each one. The equation used to compute the prediction is the following:

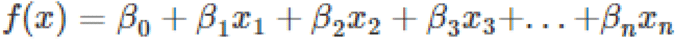

Where βn are the coefficients (being β_0_ the intercept) determined by the model and x_n_ the variables (signal values associated to markers). The predicted probabilities have been calculated with the sigmoid function:

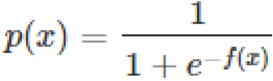

The performance of each model is internally evaluated in function of the cutoff (*p(x*) value above which an observation is positively classified) and an optimal cutoff is finally returned (cutoff at which occurs the least possible error of classification on the training dataset observations)

### Preprocessing the test datasets

As independent validation to test the selected combination and models we used different single-cell RNA sequencing datasets to see if the obtained models were able to correctly identify NK cells, without having to rely on the original 30-genes NK-signature. The test dataset were installed and/or loaded as detailed in the “Data Availability” section of this document. Both test datasets underwent the very same steps of the training dataset preparation (transposition, scaling values from 0 to 10, genes subsetting to the alphabetically ordered 30 marker genes and addition of ‘ID’ column) with the exception of the addition of ‘Class’ column which, obviously, was subsequently inferred in the end of the analysis by fitting the previously mentioned models.

### Combi-score

For each cell of the test dataset was computed a “combi-score” value (basically *p(x)*) using the standard *stats::predict()* method, specifying *type=’response’*. The combi score is, for each combination, the probability of the prediction of GLM fits on the scale of the response variable. This score was then used to assess the presence of cells classified as ‘NK’ in NK cells clusters: in this context, the combi-score is the probability of being a NK-cell given by a specific marker combination.

### Validation tests on unlabeled data

Test datasets were labeled by fitting each computed model with the *combi_score()* function following this logic:

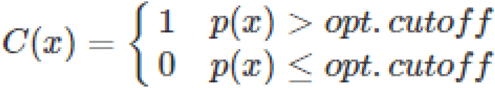

- Cells with *p(x)* higher than the optimal cutoff are classified as “NK” (= 1).
- Cells with *p(x)* lower or equal to the optimal cutoff are classified as “Other” (= 0)

The performances of classification of each combination model were obtained by comparing the inferred labels with the original cluster labels.

### Gene signature score

For each cell of the test dataset was also computed a “gene signature score” to check the effect of using selected combinations on a different published score developed for whole genes signatures. The gene signature score is described in Della Chiara et al 2021. It takes into account both the expression level and co-expression of genes within each single cell. Given a geneset, the increase of gene-signature-score is directly proportional to the number of expressed genes in the signature *and* to the sum of their level of expression. We reproduced the score computation with a custom R function described in signature_score.R script available in the GitHub combiroc package repository: https://github.com/ingmbioinfo/combiroc/blob/master/inst/external_code/signature_score.R.

### Reagents and cells

Phorbol myristate acetate (PMA), ionomycin, monensin and Brefeldin A (BFA) were purchased from SIGMA-Aldrich. The K562 cell line, was maintained in RPMI 1640 (Thermo Fisher Scientific) containing 10% FBS (GIBCO), 2 mM l-glutamine (GIBCO), 50 IU/ml penicillin (GIBCO)

### Cell isolation and purifications

Buffy coats were obtained from anonymous healthy donors from Centro Trasfusionale, Ospedale Maggiore Policlinico Milano. Human peripheral blood mononuclear cells (PBMCs) were purified through density gradient centrifugation (Ficoll-Paque Plus; GE Healthcare)

### Flow Cytometry

For flow cytometry analysis, 2×105 PBMC were stained and analyzed on FACSCANTO II (BD Bioscience). The following antibodies were used: CD56, CD3, CD122, CD94, CD16 as listed in table below. Briefly, cells were washed with MACS buffer (Miltenyi Biotech) and incubated with the antibody mix in MACS buffer at 37°C for 20 min. Then the cells were washed and analyzed on an average of 105 cells were acquired per sample, and data were analyzed using FlowJo X software (FlowJo, LLC). For CD107 degranulation assay, PBMC were resuspended at 106 cells/ml in complete RPMI 1640. Cells were then stimulated with MHC devoid, K562 cells, at an effector to target ratio of 10:1 in 96 wells plate at 200*µ*l final volume. Medium alone served as the negative control. Cells were stimulated with phorbol-12-myristate-13-acetate (PMA) (2.5 *µ*g/ml) and ionomycin (0.5 *µ*g/ml) (Sigma) as a positive control. Anti-CD107a antibody was added directly to the well at 1*µ*g. Cells were incubated for 1 h at 37 °C in 5% CO2 after which brefeldin A (Sigma) was added at a final concentration of 10 *µ*g/mL as well as 6 *µ*l of monensin (Golgi-Stop, BD Biosciences) at a final concentration of 6 *µ*g/mL and incubated for an additional 5 h at 37 8C in 5% CO2. Cells were stained for surface NK cell markers for 20 min, were then fixed with 2%PFA for 15 min at room temperature, washed with PBS and permeabilized with 0.5% Saponin and stained for intracellular tumor necrosis factor-α (TNF-α) for an additional 30 min. After washing, cells were resuspended in MACS buffer and acquired on FACSCANTO II (BD) and analyzed using FlowJo X software (FlowJo, LLC).

## List of autoantibodies used in this study

| Marker | Clone | Catalog n° | Brand |
| --- | --- | --- | --- |
| FITC Mouse Anti-Human CD3 | HIT3a | 555332 | BD Pharmingen™ |
| PE-Cy5 Mouse Anti-Human CD16 | 3G8 | 555408 | BD Pharmingen™ |
| PE-Cy7 Mouse Anti-Human CD56 | B159 | 560916 | BD Pharmingen™ |
| APC Mouse Anti-Human CD94 | HP-3D9 | 559876 | BD Pharmingen™ |
| PE Mouse Anti-Human CD107a | H4A3 | 560948 | BD Pharmingen™ |
| BV510 Mouse Anti-Human CD122 | Mik-β3 | 563093 | BD Horizon™ |
| BV421 Mouse Anti-Human TNF-α | MAb11 | 566275 | BD Horizon™ |
| APC-Cy7 Mouse Anti-Human IFN-γ | 502529 | 502529 | BioLegend |

## Acknowledgements

We would like to thank Jens Geginat, Davide Cittaro, Asia Zonca and Stefano Biffo for fruitful discussion and comments, members of the Bioinformatics Units at INGM for constructing criticism. S.M. was funded by the European Research Council (ERC) Consolidator Award MicroC 772970 (to PI F. Buffa).

## Author contributions

R.L.R. proposed the package project and coordinated the whole work; I.F. conceived new features; R.L.R, M.B, I.F. designed the analyses; I.F. and R.L.R. performed analyses, wrote code, compiled the package, wrote manuscript, produced figures and vignettes; S.M. drafted code parts; M.B. searched and selected datasets; R.L.R., S.M. reviewed and debugged code; M.C., M.B. performed cytometric and cytotoxicity measurements and analyzed assays’ data; S.A., R.G., secured funding; all authors reviewed the manuscript.

### Data and Code availability

All data used in this paper is already available for download from respective publication sources. Details on their usage and loading in the combiroc single cell workflow can be found in the accompanying vignette https://ingmbioinfo.github.io/combiroc/articles/combiroc_vignette_2.html

- Multimodal PBMC dataset (training dataset) from Hao et al. 2021 can be downloaded as an h5 Seurat data (h5s) from the Fred Hutchinson & New York Genome Center Atlas page (https://atlas.fredhutch.org/nygc/multimodal-pbmc/)
- CBMC-CITE-seq dataset (testing dataset) shown in ‘Using Seurat with multimodal data’ vignette, can be directly loaded from the SeuratData package.
- PBMC3K (testing dataset): installed from the SeuratData library: https://github.com/satijalab/seurat-data.
- COVID-19 PBMC (testing dataset) Ncl-Cambridge-UCL from Haniffa lab : https://www.covid19cellatlas.org/index.patient.html.

The combiroc package in the most recent development version is available from our GitHub repository (https://ingmbioinfo.github.io/combiroc/) as well as from the Comprehensive R Archive Network (https://cran.r-project.org/) as a stable version. The whole workflow with R code, datasets, precomputed objects, and commented results are available for review and reproducibility in the mentioned GitHub repository in the dedicated pages

- Supplementary materials 1: Standard workflow for the combiroc package: “Guide to CombiROC package - Standard workflow” on GitHub at https://ingmbioinfo.github.io/combiroc/articles/combiroc_vignette_1.html
- Supplementary materials 2: Single cell sequencing complete workflow: “Signature refining tutorial - scRNA-seq Workflow” Available on GitHub at https://ingmbioinfo.github.io/combiroc/articles/combiroc_vignette_2.html
- Supplementary materials 3: Zipped folder containing scripts and results of analyses performed on the PBMC-Covid19 (testing) dataset.

## References

1. Mazzara, S. et al. CombiROC: an interactive web tool for selecting accurate marker combinations of omics data. Sci. Rep. 7, 45477 (2017).

2. Bombaci, M. & Rossi, R. L. Computation and selection of optimal biomarker combinations by integrative ROC analysis using combiroc. Methods Mol. Biol. 1959, 247–259 (2019).

3. Bombaci, M. et al. Novel biomarkers for primary biliary cholangitis to improve diagnosis and understand underlying regulatory mechanisms. Liver Int. 39, 2124–2135 (2019).

4. Sola, L. et al. Enhancing antibody serodiagnosis using a controlled peptide coimmobilization strategy. ACS Infect. Dis. 4, 998–1006 (2018).

5. CRAN - Package combiroc. https://CRAN.R-project.org/package=combiroc.

6. Abdelaal, T. et al. A comparison of automatic cell identification methods for single-cell RNA sequencing data. Genome Biol. 20, 194 (2019).

7. Grabski, I. N. & Irizarry, R. A. A probabilistic gene expression barcode for annotation of cell types from single-cell RNA-seq data. Biostatistics 23, 1150–1164 (2022).

8. Narni-Mancinelli, E., Vivier, E. & Kerdiles, Y. M. The “T-cell-ness” of NK cells: unexpected similarities between NK cells and T cells. Int. Immunol. 23, 427–431 (2011).

9. Hao, Y. et al. Integrated analysis of multimodal single-cell data. Cell 184, 3573-3587.e29 (2021).

10. Stoeckius, M. et al. Simultaneous epitope and transcriptome measurement in single cells. Nat. Methods 14, 865–868 (2017).

11. Satija, R., Farrell, J. A., Gennert, D., Schier, A. F. & Regev, A. Spatial reconstruction of single-cell gene expression data. Nat. Biotechnol. 33, 495–502 (2015).

12. Stephenson, E. et al. Single-cell multi-omics analysis of the immune response in COVID-19. Nat. Med. (2021) doi:10.1038/s41591-021-01329-2.

13. Mariotti, F. R., Quatrini, L., Munari, E., Vacca, P. & Moretta, L. Innate Lymphoid Cells: Expression of PD-1 and Other Checkpoints in Normal and Pathological Conditions. Front. Immunol. 10, 910 (2019).

14. Della Chiara, G. et al. Epigenomic landscape of human colorectal cancer unveils an aberrant core of pan-cancer enhancers orchestrated by YAP/TAZ. Nat. Commun. 12, 2340 (2021).

15. Uhlén, M. et al. Tissue-based map of the human proteome. Science 347, 1260419 (2015).

16. Wu, Y., Tian, Z. & Wei, H. Developmental and functional control of natural killer cells by cytokines. Front. Immunol. 8, 930 (2017).

