## Supplemental text and legends for "Combiroc: when ‘less is more’ in bulk and single cell marker signatures"

#### Supplementary Figure legends

**Suppl. Figure 1 - a.** UMAP visualization of the annotated cell clusters of multimodal-PBMC CITE-seq atlas from Hao et al. 2021. Cell clusters are annotated for eight different cell types according to gene expression and CITE-seq results in the original work. **b.** Feature plots of the top four differentially NK cell markers belonging to the NK specific gene marker signature: most of them are also highly expressed in other non-NK clusters (e.g. CD8 T), making them too unspecific to allow a well-defined cluster annotation when considered individually.

**Suppl. Figure 2 - a.** ROC curves for the top four individual NK markers and for the top four marker combinations determined with combiroc. **b.** Table of performance parameters for the markers (feature) depicted: AUC, area under the curve; SE, sensitivity; SP, specificity; C-Off, optimal cut off; ACC, accuracy; TN, true negative; TP, true positive; FN, false negative; FP, false positive; NPV, negative predictive value; PPV, positive predictive value.

**Suppl. Figure 3 -** Cell clusters UMAP visualizations and violin plots for cluster specific expression of the top four individual gene markers in three testing datasets: (**a**, **b**) CBMC dataset, (**c**, **d**) PBMC-3K dataset and (**e**, **f**) PBMC-Covid19 dataset.

**Suppl. Figure 4 - a**, Gating strategy to discriminate among the different NK cells subsets. Live lymphocytes were gated based on side and forward scatter dot plot display during the acquisition process. The CD3- lymphocyte population was identified using a negative gating strategy. **b**, CD107a expression is highly elevated on the surface of NK cells following stimulation. Flow cytometry analysis reveals the percentage of CD3-/CD56+ and CD3-/CD122+ cells expressing CD107a after stimulation with PMA/ionomycin and with K562 cells. The presented percentages are the results of four independent experiments. Fractions of NK cells identified with FACS analysis as CD56+/CD16+ cells and as CD122+/CD94+ cells are shown in the two upper right panels.

### Supplementary Table

**Suppl. Table 1 -** Enrichment of marker genes belonging to the selected combinations in "Blood and Immune" cell types from the single cell data of Human Protein Atlas (HPA, https://www.proteinatlas.org/). Rightmost column reports the most enriched cell types from single cell in HPA and the relative specificity score.

| Gene | Aliases | Description | Localization | Most enriched (specificity score) |
|---|---|---|---|---|
| IL2RB | CD122 | Interleukin 2 receptor subunit beta | *Membrane* | NK cells, T cells (**0.89**) |
| KLRF1 | CLEC5C | Killer Cell Lectin Like Receptor F1 | *Membrane* | NK cells (**0.90**) |
| KLRD1 | CD94 | Killer Cell Lectin Like Receptor D1 | *Membrane* | NK cells (**0.83**) |
| SPON2 | DIL1 | Spondin 2 | Intracellular | NK cells, dendritic cells (**0.66**) |
| TRDC1 | TCRD | T Cell Receptor Delta Constant | *Membrane* | NK cells, T cells (**0.93**) |
| GNLY | TLA519 | Granulysin | Intracellular | NK cells (**0.84**) |
| FCGR3A | CD16 | Fc Gamma Receptor IIIa | *Membrane* | Monocytes, NK-cells, Macrophages, Hofbauer cells, Kupffer cells (**0.84**) |
| MYOM2 | TTNAP | Myomesin 2 | Intracellular | NK-cells (**0.76**) |
| GZMB | | Granzyme B | Intracellular | Dendritic cells, NK-cells (**0.89**) |
| CLIC3 | | Chloride Intracellular Channel 3 | Intracellular | Dendritic cells, NK-cells (**0.70**) |

### Supplementary material

**Suppl. Material 1 -** Standard workflow for the combiroc package: "Guide to CombiROC package - Standard workflow" on GitHub at https://ingmbioinfo.github.io/combiroc/articles/combiroc_vignette_1.html

**Suppl. Material 2 -** Single cell sequencing complete workflow: "Signature refining tutorial - scRNA-seq Workflow" Available on GitHub at https://ingmbioinfo.github.io/combiroc/articles/combiroc_vignette_2.html

**Suppl. Material 3 –** Scripts and results of analyses performed on the PBMC-Covid19 (testing) dataset were not organized for display in a vignette but are available as a Zip archive upon request.
