## Supplementary figures and images for "Combiroc: when ‘less is more’ in bulk and single cell marker signatures"

### Supplemental Figure 1

**a**

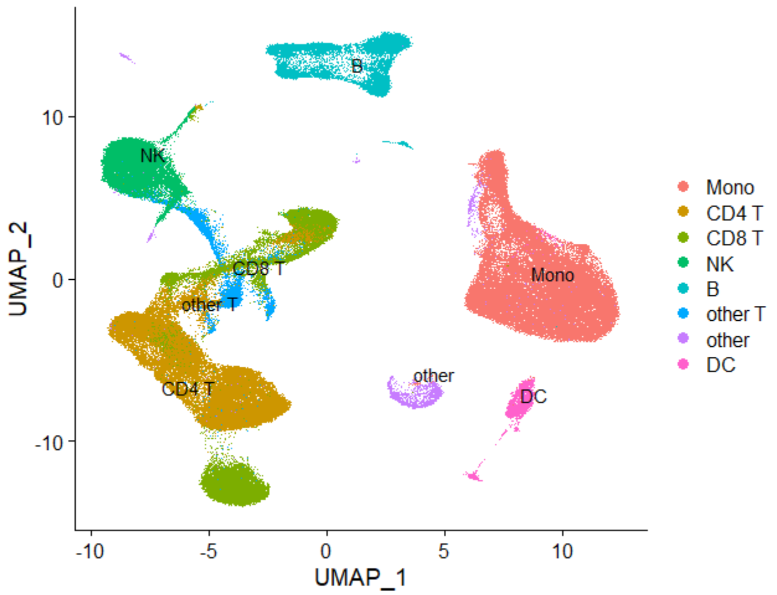


**b**

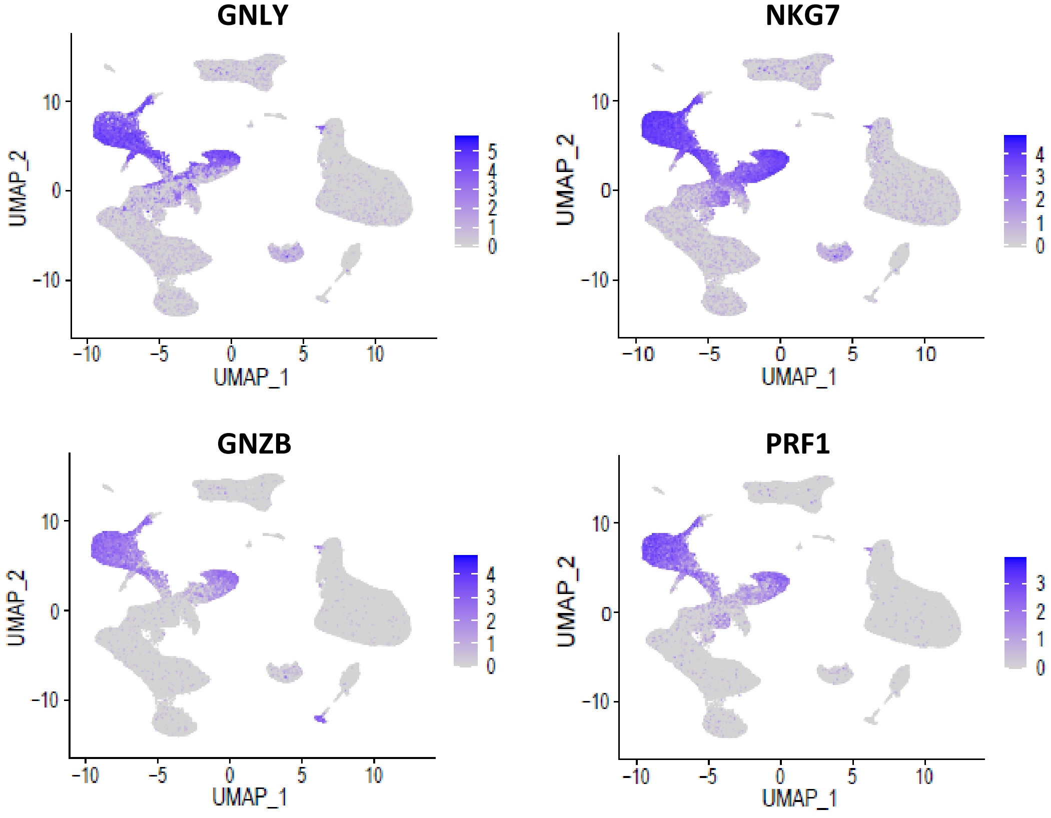


#### Supplementary Figure 1

### Supplemental Figure 3_1

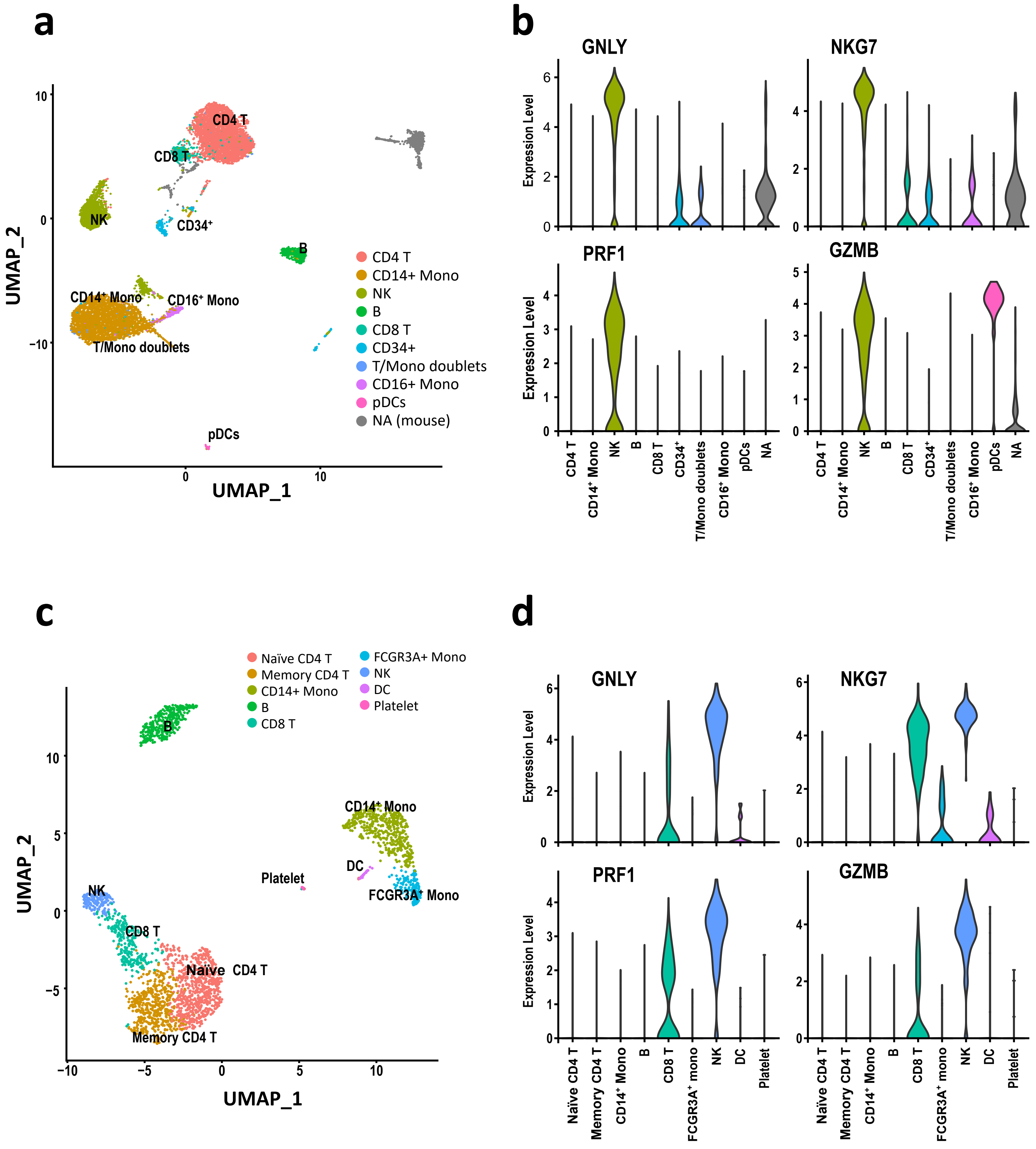


#### Supplementary Figure 3

### Supplemental Figure 3_2

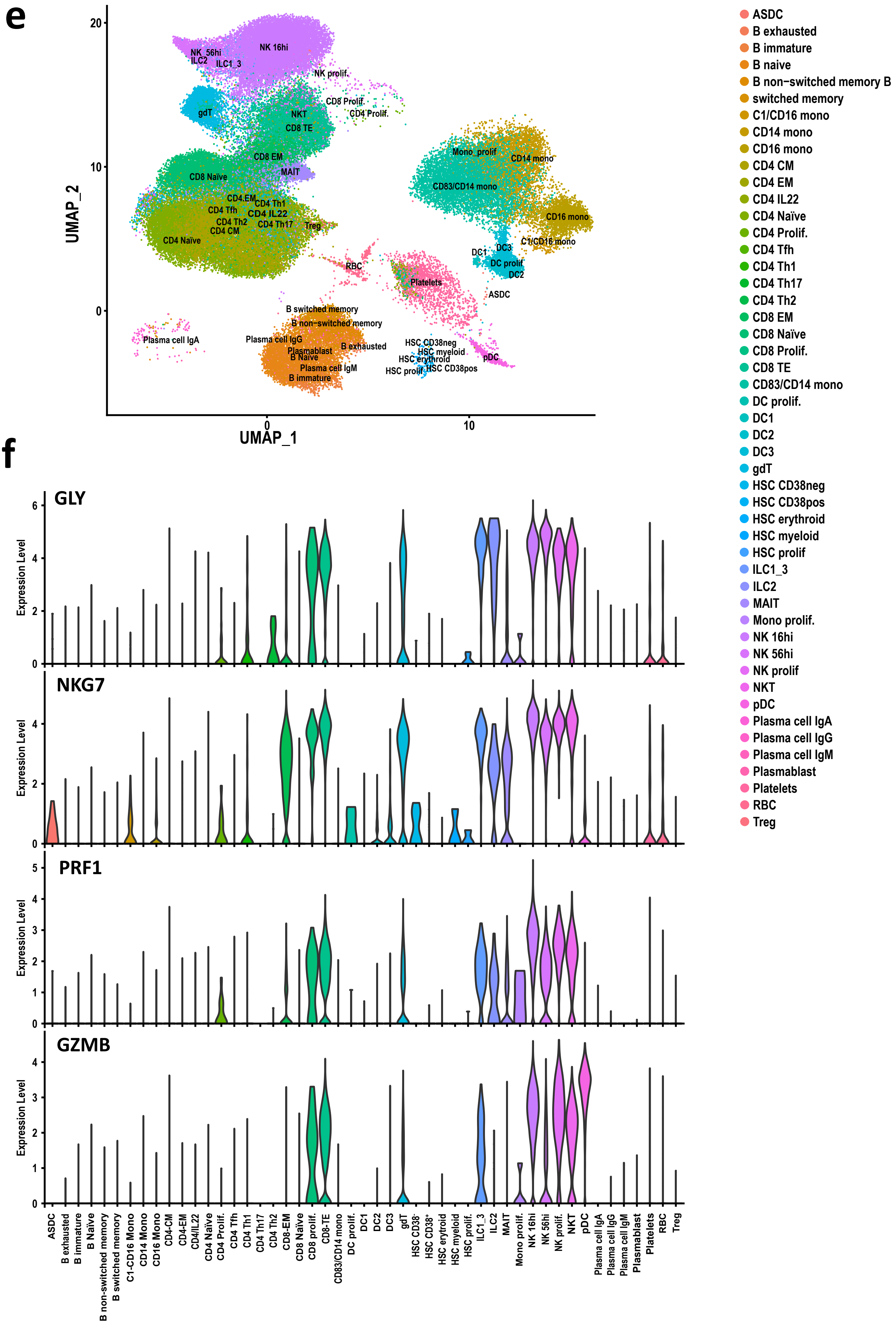


**Supplementary Figure 3** - *cont.*

### Supplemental Figure 4

**a**

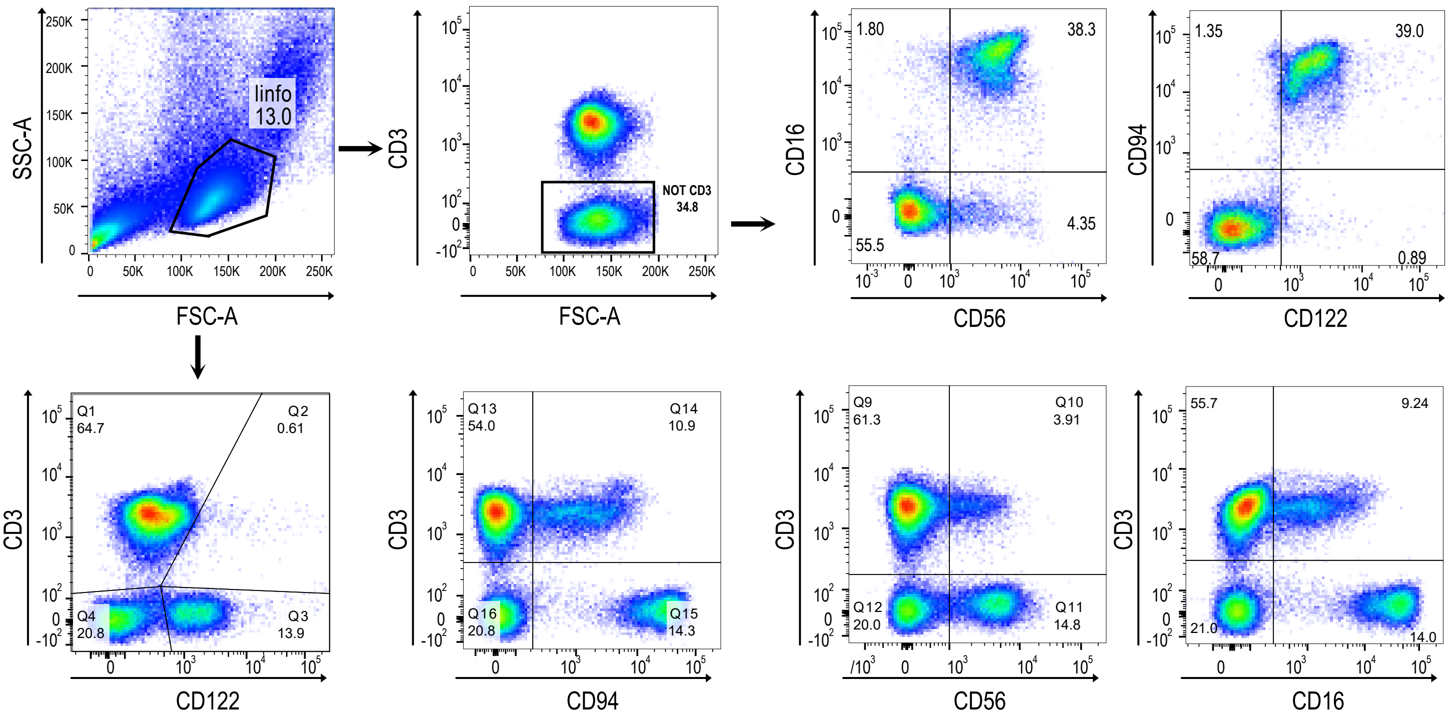


**b**

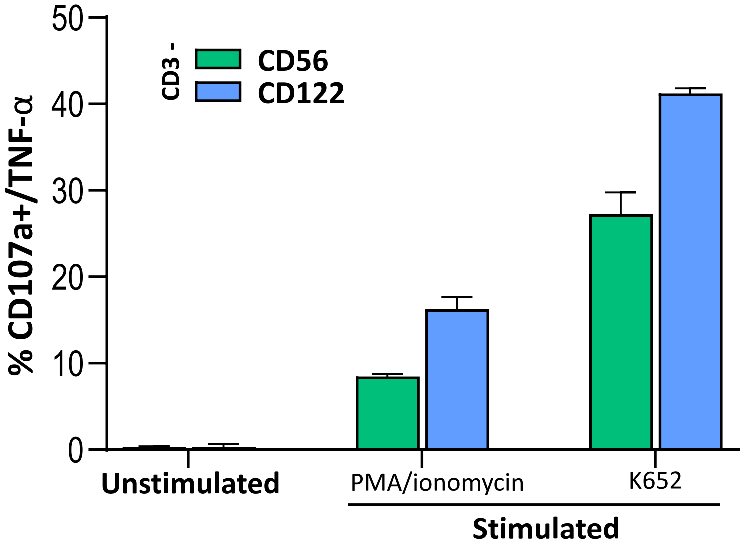


#### Supplementary Figure 4
