## Supplemental Figure 2 for "Combiroc: when ‘less is more’ in bulk and single cell marker signatures"

**a**

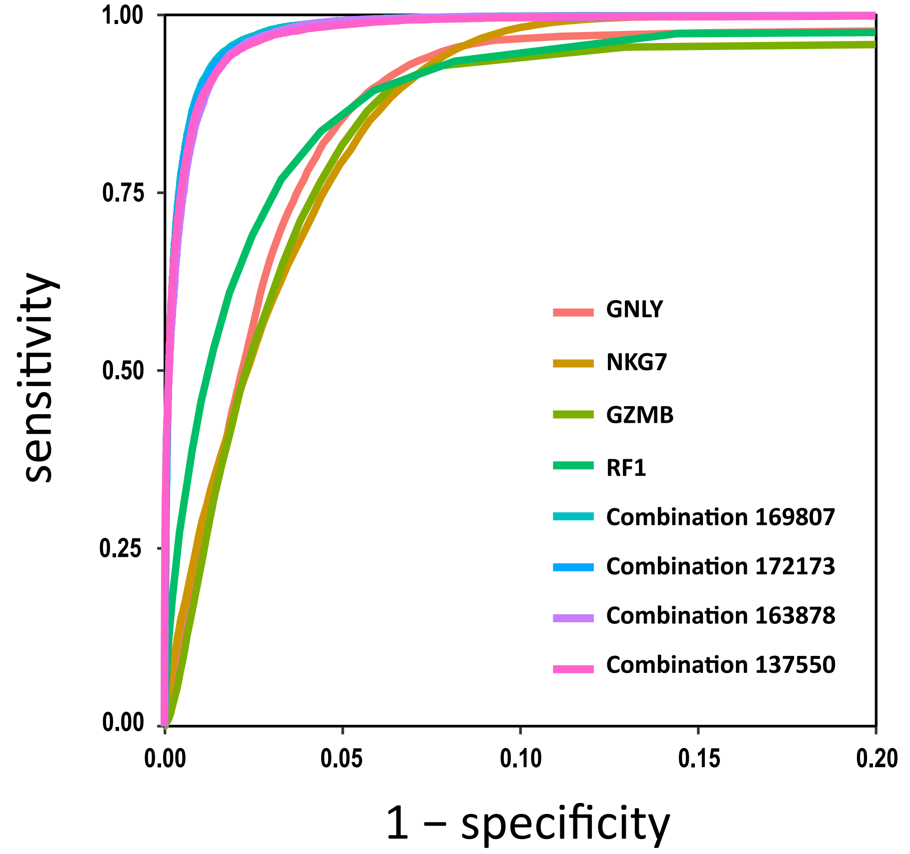


**b**

| Feature | AUC | SE | SP | C-Off | ACC | TN | TP | FN | FP | NPV | PPV |
|---|---|---|---|---|---|---|---|---|---|---|---|
| **GNLY** | 0.962 | 0.959 | 0.914 | 0.146 | 0.92 | 130861 | 17908 | 756 | 12239 | 0.994 | 0.594 |
| **NKG7** | 0.970 | 0.982 | 0.901 | 0.131 | 0.911 | 128972 | 18336 | 328 | 14128 | 0.997 | 0.565 |
| **GZMB** | 0.949 | 0.928 | 0.925 | 0.136 | 0.925 | 132375 | 17316 | 1348 | 10725 | 0.990 | 0.618 |
| **PRF1** | 0.965 | 0.935 | 0.919 | 0.137 | 0.921 | 131569 | 17452 | 1212 | 11531 | 0.991 | 0.602 |
| **Combination 169807** | 0.995 | 0.982 | 0.968 | 0.107 | 0.97 | 138553 | 18326 | 338 | 4547 | 0.998 | 0.801 |
| **Combination 172173** | 0.995 | 0.977 | 0.970 | 0.108 | 0.971 | 138769 | 18229 | 435 | 4331 | 0.997 | 0.808 |
| **Combination 163878** | 0.995 | 0.985 | 0.962 | 0.078 | 0.965 | 137680 | 18393 | 271 | 5420 | 0.998 | 0.772 |
| **Combination 137550** | 0.995 | 0.976 | 0.966 | 0.101 | 0.967 | 138179 | 18224 | 440 | 4921 | 0.997 | 0.787 |

### Supplementary Figure 2
